# Inhibition of the melanocortin-3 receptor (MC3R) causes generalized sensitization to anorectic agents

**DOI:** 10.1101/2023.12.05.570114

**Authors:** Naima S. Dahir, Yijun Gui, Yanan Wu, Patrick R. Sweeney, Savannah Y. Williams, Luis E. Gimenez, Tomi K. Sawyer, Stephen T. Joy, Anna K. Mapp, Roger D. Cone

**Author notes:** Corresponding Author: Roger D. Cone, Ph.D. Life Sciences Institute, University of Michigan 210 Washtenaw Ave. Ann Arbor, MI 48109-2216. Conflict of Interest Statement: RDC, PS, SYW, TS, and the University of Michigan are shareholders in Courage Therapeutics. RDC, NSD, PS, and TS are on patents related to this work.

## Abstract

The melanocortin-3 receptor (MC3R) acts presynaptically to regulate GABA release from agouti-related protein (AgRP) nerve terminals and thus may be a negative regulator of multiple circuits involved in feeding behavior and energy homeostasis. Here, we examined the role of MC3R in regulating the response to various anorexigenic agents. Our findings reveal that genetic deletion or pharmacological inhibition of MC3R improves the dose responsiveness to Glucagon-like peptide 1 (GLP1) agonists, as assayed by inhibition of food intake and weight loss. An enhanced anorectic response to other agents, including the acute satiety factors peptide YY (PYY_3-36_) and cholecystokinin (CCK) and the long-term adipostatic factor, leptin, demonstrated that increased sensitivity to anorectic agents is a generalized result of MC3R antagonism. Enhanced neuronal activation in multiple nuclei, including ARH, VMH, and DMH, was observed using Fos immunohistochemistry following low-dose liraglutide in MC3R knockout mice (*Mc3r^-/-^*), supporting the hypothesis that the MC3R is a negative regulator of circuits regulating multiple aspects of feeding behavior. The enhanced anorectic response in *Mc3r ^-/-^* mice after administration of GLP1 analogs was also independent of the incretin effects and malaise induced by GLP1R analogs, suggesting that MC3R antagonists may have value in enhancing the dose-response range of obesity therapeutics.

## Introduction

The central melanocortin system is fundamental in regulating feeding behavior and energy homeostasis. Mutations in the melanocortin-4 receptor (MC4R) or genetic deletion of *Mc4r* results in hyperphagia, hyperinsulinemia, and obesity^1–3^. In contrast, *Mc3r* deletion does not produce hyperphagia or hyperinsulinemia in ad libitum-fed animals^4,5^. However, the MC3R acts as a negative regulator of the MC4R system, controlling responses of the melanocortin system to both internal and external challenges^6–10^, controlling the upper and lower boundaries of energy homeostasis^9^ and providing sexually dimorphic inputs to the system^8,10,11^. For example, MC3R is essential in promoting normal compensatory re-feeding and neuroendocrine responses to fasting^7–10^. MC3R accomplishes this by regulating GABA release onto MC4R neurons from presynaptic MC3R expressed on AgRP neurons^9,12^, and by controlling the activation of AgRP neurons^13^. Further, MC3R is requisite for normal growth^5,14^ and reproductive function, with mutations leading to reduced linear growth and lean mass and late onset of puberty in mice and humans^15^.

We have previously examined the effect of the MC3R on MC4R-expressing paraventricular hypothalamic (PVH) neurons and demonstrated that MC3R blockade enhances the magnitude and duration of the inhibition of food intake in response to a single dose of the MC4R agonist LY2112688^9^. MC3R is expressed in nearly all AgRP neurons, and the AgRP neurons are known to broadly impact multiple aspects of feeding behavior and suppress competing behavioral states, including fear, alarm, and thirst. For example, we have demonstrated that deletion of the MC3R increases sensitivity to anorexia induced by restraint and social isolation^10^. Thus, MC3R activity may be found to regulate responsiveness to anorectic stimuli broadly. Here, we examined the ability of the MC3R to regulate responsiveness to a variety of anorexigenic agents, from acute satiety factors like CCK and PYY_3-36_ to the long-term adipostatic factor leptin, as well as to leading GLP1-based therapeutic agents like liraglutide and tirzepatide.

## Results

### Disruption of MC3R signaling increases sensitivity to GLP1R agonists

Previously, we reported that deletion of MC3R (*Mc3r* ^-/-^) in male and female mice resulted in enhanced acute inhibition of food intake in response to a single dose of the glucagon-like peptide-1 receptor agonist (GLP1R), liraglutide^10^ We tested the effect of various concentrations of liraglutide in mice lacking MC3R and their wild-type counterparts. In this testing paradigm, we administered concentrations of 0.05 mg/kg – 0.4 mg/kg of liraglutide at least 30 minutes before the onset of the dark cycle. We manually measured 24-hour food intake (spillage included) and body weight change in male and female mice. As expected, liraglutide reduced food intake and body weight in wild-type male (Figure 1A-B) and female (Figure 1C-D) mice in a concentration-dependent manner. In the *Mc3r ^-/-^*, we found an increased dose sensitivity to liraglutide-induced suppression of food intake and weight loss at 24 h post-administration. Unlike wild-type mice, *Mc3r ^-/-^* mice exhibited reduced food intake and weight loss in response to low concentrations of liraglutide, 0.05mg/kg, indicating hypersensitivity to the compound. Next, we wanted to test the sensitivity of *Mc3r^-/-^* mice to other GLP1R agonists. We tested the impact of semaglutide, a secondary GLP1 drug shown to have a longer half-life than liraglutide, thereby increasing both the incretin and anorectic duration of drug action^16^. We tested male mice and found that, like liraglutide, deletion of MC3R leads to increased responsiveness to semaglutide; *Mc3r^-/-^* mice responded more to both the anorectic effects and weight loss in response to semaglutide than the wildtype (WT) mice, illustrating that this hypersensitivity was seen in response to multiple GLP1R agonists (Figure 1SA-B).

**Figure 1.**
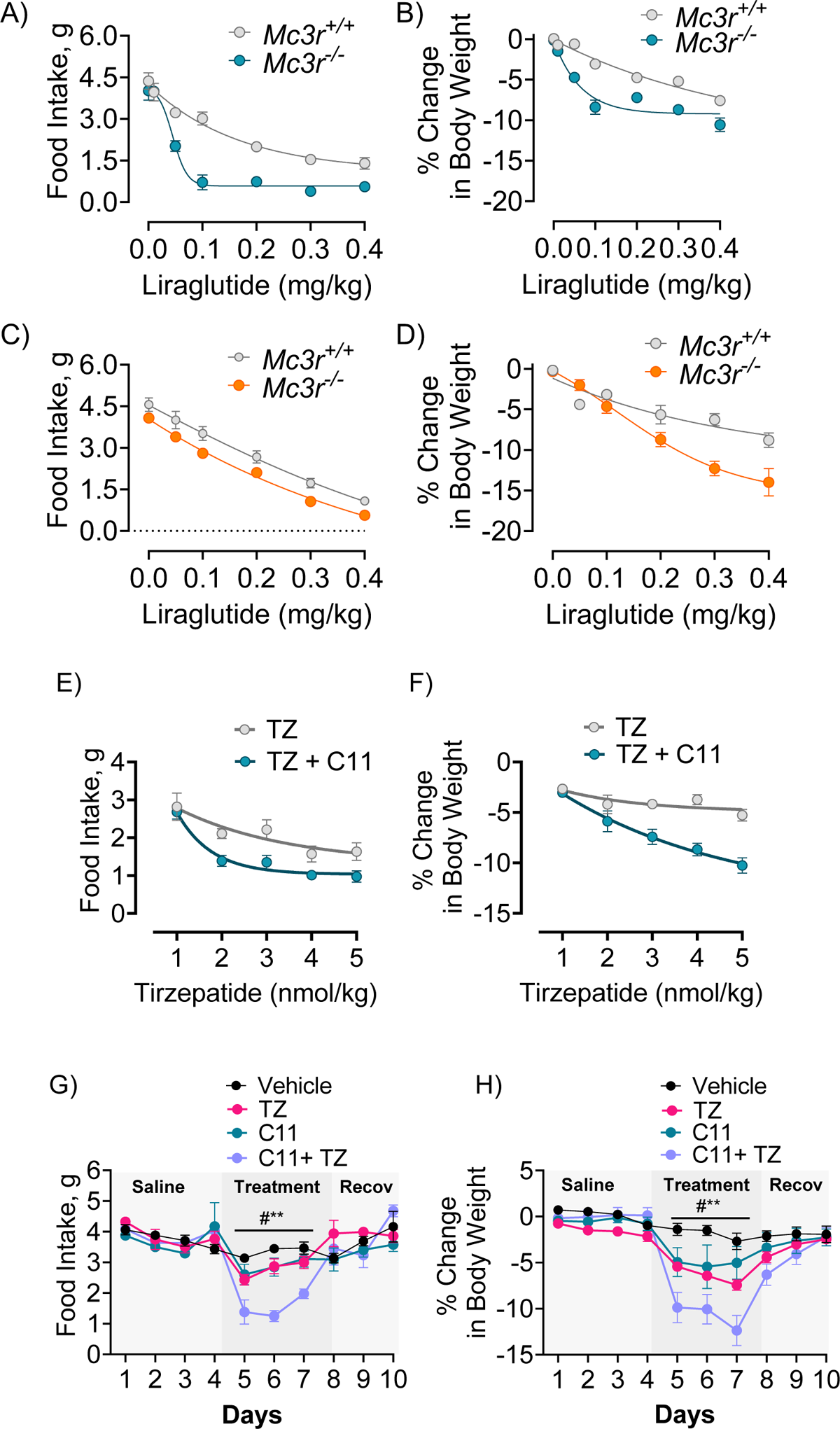
Loss of MC3R increases responsiveness to GLP1 drugs. Liraglutide (0.05mg/kg – 0.4mg/kg) administration results in more significant inhibition of (A) food intake and (B) weight loss in *Mc3r* ^-/-^ male mice compared to *Mc3r ^+/+^* mice in a dose-dependent manner after 24-hours (N = 7–8/group). (C) Liraglutide-induced feeding (D) and body weight changes of *Mc3r ^+/+^*and *Mc3r ^-/-^* female mice (n = 7–8/group; 0.05mg/kg – 0.4mg/kg). Tirzepatide (TZ, 1nmol/kg – 4nmol/kg) and coadministration of tirzepatide and C11-induced (E) feeding responses and (F) body weight changes of wildtype male mice (vehicle, n = 10, all other groups n = 8) at 24-hours post-injection. (G) 24-hour feeding and (H) body weight change after chronic injections of tirzepatide (2nmol/kg), C11 (0.5nmol), tirzepatide and C11, and vehicle. Data is expressed as mean ± SEM, and statistical tests were two-way ANOVA with Tukey’s test for post hoc analysis for (G-J). For all the dose-response curve data, repeated measures of two-way ANOVA were corrected for multiple comparisons using the Tukey–Kramer method for each time point, and data were fitted with four parameters: nonlinear fit, *p < 0.05, *p <0.01, ***p <0.001.

We have previously shown that MC3R peptide antagonist compound 11 (C11)^17^ can reliably inhibit the activity of the MC3R and co-administration of C11 and liraglutide can further lower the body weight of wildtype mice more than liraglutide alone^10^. We repeated the same experiment and additionally used *Mc3r^-/-^*mice to illustrate the specificity of C11. We found that pharmacological inhibition of MC3R (C11; 1nmol/1µl, intracerebroventricularly, icv) and liraglutide, together, resulted in profound weight loss and decreased food intake over 8-hr, 12-hr, and 24-hr periods (Figure 2S). Furthermore, no effect was seen in mice with MC3R deletion, validating the pharmacological specificity of C11.

**Figure 2.**
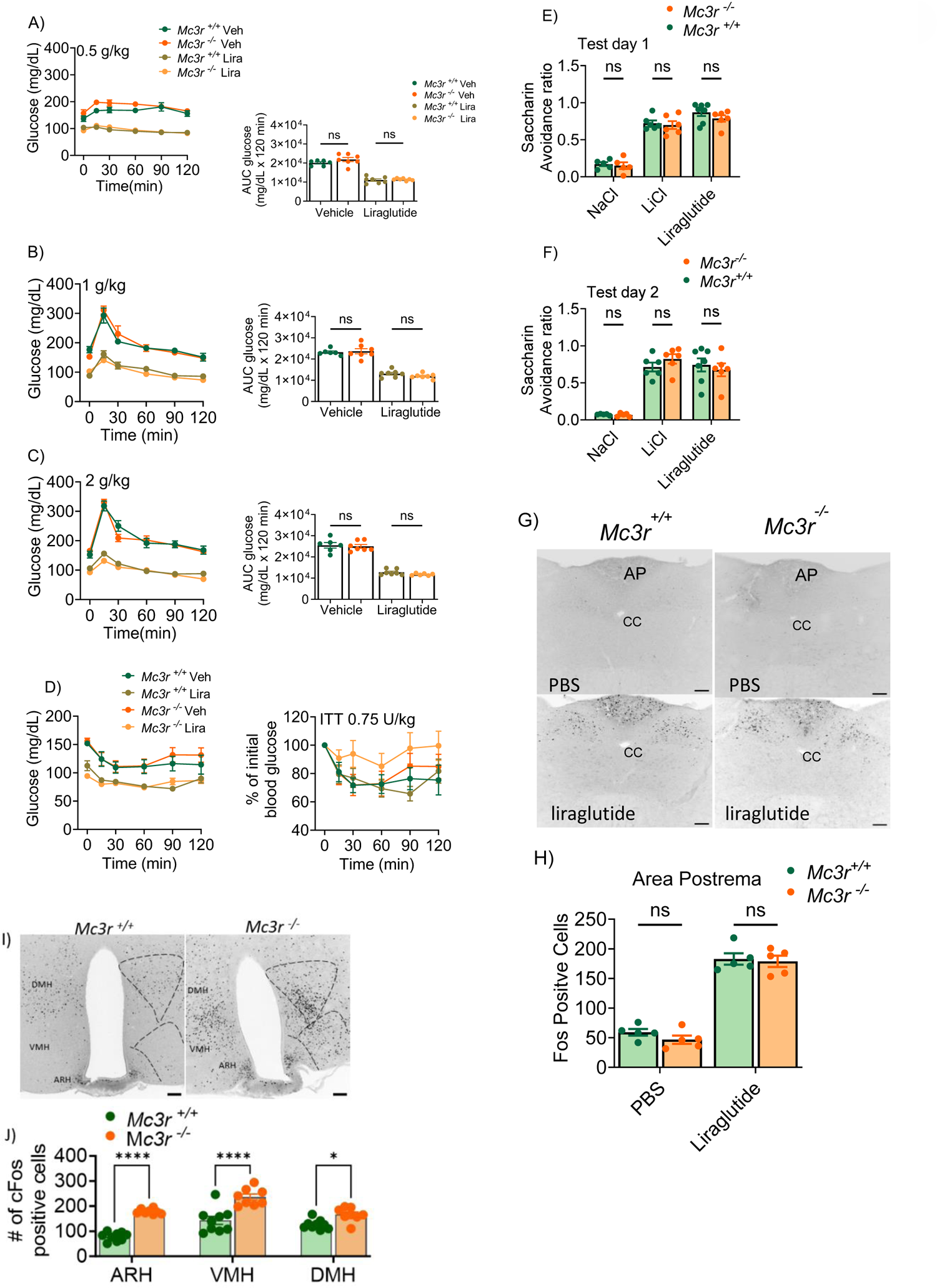
MC3R deletion has no effect on the incretin activity or malaise associated with liraglutide. Glucose levels and area under the curve before and after oral administration of glucose at 0.5 g/kg (A), 1.0 g/kg (B), 2.0 g/kg (C) after liraglutide (0.2 mg/kg), and vehicle treatment in *Mc3r^+/+^* and *Mc3r^-/--^* male mice (n = 7/group). (D) Glucose levels before and after insulin injection (0.75 IU insulin/kg) in *Mc3r^+/+^* and *Mc3r^-/-^*male mice treated with liraglutide and vehicle (n = 8/group). CTA test days 1 (E) and 2 (F) after liraglutide administration in *Mc3r^+/+^* and *Mc3r^-/-^* male mice. (G) Representative images from the AP showing Fos IHC after saline or liraglutide injection were obtained from *Mc3r^+/+^* and *Mc3r^-/-^* male mice. (H) quantified cells expressing Fos after saline or liraglutide injection in *Mc3r*^+/+^ and *Mc3r^-/-^* male mice. (I) Representative images from the hypothalamus showing Fos IHC after liraglutide treatment in *Mc3r^+/+^*and *Mc3r^-/-^* mice. (J) quantified cells expressing Fos in ARH, VMH, and DMH after liraglutide treatment.

Next, we examined the weight loss and anorectic effects of MC3R pharmacological inhibition in response to increasing doses of tirzepatide. This diabetes drug acts as a dual agonist for GLP1R and the glucose-dependent insulinotropic polypeptide (GIP) in humans^18,19^. Administration of tirzepatide (1nmol/kg - 5nmol/kg, sc) in wild-type mice decreased food intake and promoted weight loss in male mice (Figure 1E-F) measured 24 hours following administration. Furthermore, co-administration of C11 (1nmol/1µl, icv) and varying dosages of tirzepatide profoundly decreased 24-hour food intake and 24-hour body weight after a single injection of both compounds (Figure 1E-F). To determine if chronic administration of tirzepatide and C11 further promotes their hypersensitivity effects, 24-hour food intake and body weight of male mice administered with C11 (0.5nmol, icv), tirzepatide (2nmol/kg, sc), C11 and tirzepatide, and vehicle (DMSO, icv and saline, sc) were measured; 4 days of vehicle treatment, followed by treatment for 3 days, and recorded recovery for an additional 3 days are shown (Figure 1G-H). Administration of C11 and tirzepatide alone showed comparable effects on weight loss and decreased food intake compared to vehicle-treated groups. However, administering both compounds produced greater inhibition of food intake and weight loss than either compound, demonstrating higher efficacy of tirzepatide with MC3R antagonism in a chronic treatment model.

### Deletion of MC3R does not increase the incretin effects or malaise induced by liraglutide

GLP1 analogs have been historically associated with their incretin effects, stimulating insulin secretion and suppressing glucagon secretion in hyperglycemic or euglycemic states^20^. All the GLP1R analogs approved by the FDA are reliably used to treat Type II diabetes and regulate glucose homeostasis^21^. We tested whether *Mc3r* ^-/-^ mice also exhibited increased sensitivity to the incretin effects of GLP1-analogs by measuring oral glucose (GTT) and insulin (ITT) tolerance and insulin and GLP1 levels. WT and *Mc3r^-/-^* mice were fasted and injected with liraglutide (0.2mg/kg, sc) or vehicle (PBS, sc). After 6 hours of fasting, glucose responses were measured following 0.5g/kg, 1g/kg, and 2g/kg of an oral glucose bolus. However, we found no effect of genotype; both *Mc3r^+/+^* and *Mc3r^-/-^* mice responded similarly to the oral GTT (Figure 2A-C). Liraglutide consistently produced significantly better glucose disposal than a vehicle in both genotypes (Fig 2A-C). Next, we repeated the same experiment as the GTT, except we injected an insulin bolus (0.75 IU insulin/kg, i.p.) *Mc3r^+/+^* and *Mc3r^-/-^* mice and measured glucose disposal over 2 h. Similarly, we observed glucose lowering by liraglutide but no effect of genotypes (*Mc3r^+/+^* vs. *Mc3r^-/-^*) (Fig 2D).

The most reported adverse effects of GLP1R agonists are gastrointestinal; patients report increased emetic responses^22–25^, as well as exacerbating gastroparesis^26^. To test whether our observed hypersensitive anorectic responses to GLP1-analogs were accompanied by increased malaise, we tested whether liraglutide triggered increased conditioned taste aversion in *Mc3r^-/-^* mice as previously described^24,27^. Low concentrations of liraglutide that resulted in decreased food intake and promoted weight loss (0.05mg/kg) in *Mc3r^-/-^* mice were paired with the non-nutritive sweetener saccharin (0.1%). A second group of mice received saccharin paired with lithium chloride (150 mM) to induce gastric malaise. A third group received saccharin paired with saline (150 mM NaCl). Mice underwent training for two days, conditioning for two days, and testing for two days. *Mc3r^+/+^* and *Mc3r^-/-^* mice had low avoidance for saccharin when paired with saline and high avoidance when paired with either the liraglutide or LiCl. We found that genetic deletion of MC3R did not increase the conditioned taste aversion associated with administering liraglutide (Fig 2E-F). GLP1 agonists are also known to activate neurons in the area postrema at doses that induce CTA. Deletion of the MC3R was not observed to increase activation of area postrema neurons in response to 0.1mg/kg liraglutide Figure 2G-H). In contrast, a profound increase in neuronal activation following liraglutide treatment was seen in hypothalamic feeding circuits in the VMH, DMH, and ARH of the *Mc3r^-/-^*mouse relative to WT mice (Figure 2I-J). Collectively, these data illustrate that the observed anorectic hypersensitization of GLP1-analogs in *Mc3r^-/-^* mice is independent of emetic responses.

### Deletion of MC3R produces increased sensitivity to diverse anorectic hormones

We next sought to determine if the role of the MC3R in sensitivity to GLP1 agonists was unique to this family of hormones or more generalizable. We first tested the effect of the long-term adipostatic hormone leptin, a hormone produced by the adipose tissue in proportion to fat stores^28^. Given that *Mc3r^-/-^* mice display late-onset weight gain and hyperleptinemia^7,13^, these mice should theoretically exhibit leptin resistance. In contrast, *Mc3r^-/-^* mice also exhibited increased anorectic sensitivity to leptin; low doses of leptin, which otherwise did not affect food intake in WT mice, showed robust anorectic response in male *Mc3r^-/-^* mice (Fig 3A). The effect was not observed in female mice (Fig 3B), although the estrous cycle, a significant determinant of daily food intake and leptin levels^29,30^, was not synchronized in this study. We next tested gut hormones known to act acutely as satiety factors, including a form of the peptide YY (PYY_3-36_) and cholecystokinin (CCK), both secreted by enteroendocrine cells in the small intestine. We found that male and female *Mc3r^-/-^* mice had increased responses to PYY_3-36_ in a dose-dependent manner over 2-h nocturnal feeding (Fig 3C-D). Similarly, administration of CCK produced an increased anorectic response in male and female *Mc3r^-/-^* mice compared to the wildtype mice in a dose-dependent manner (Fig 3E-F).

**Figure 3.**
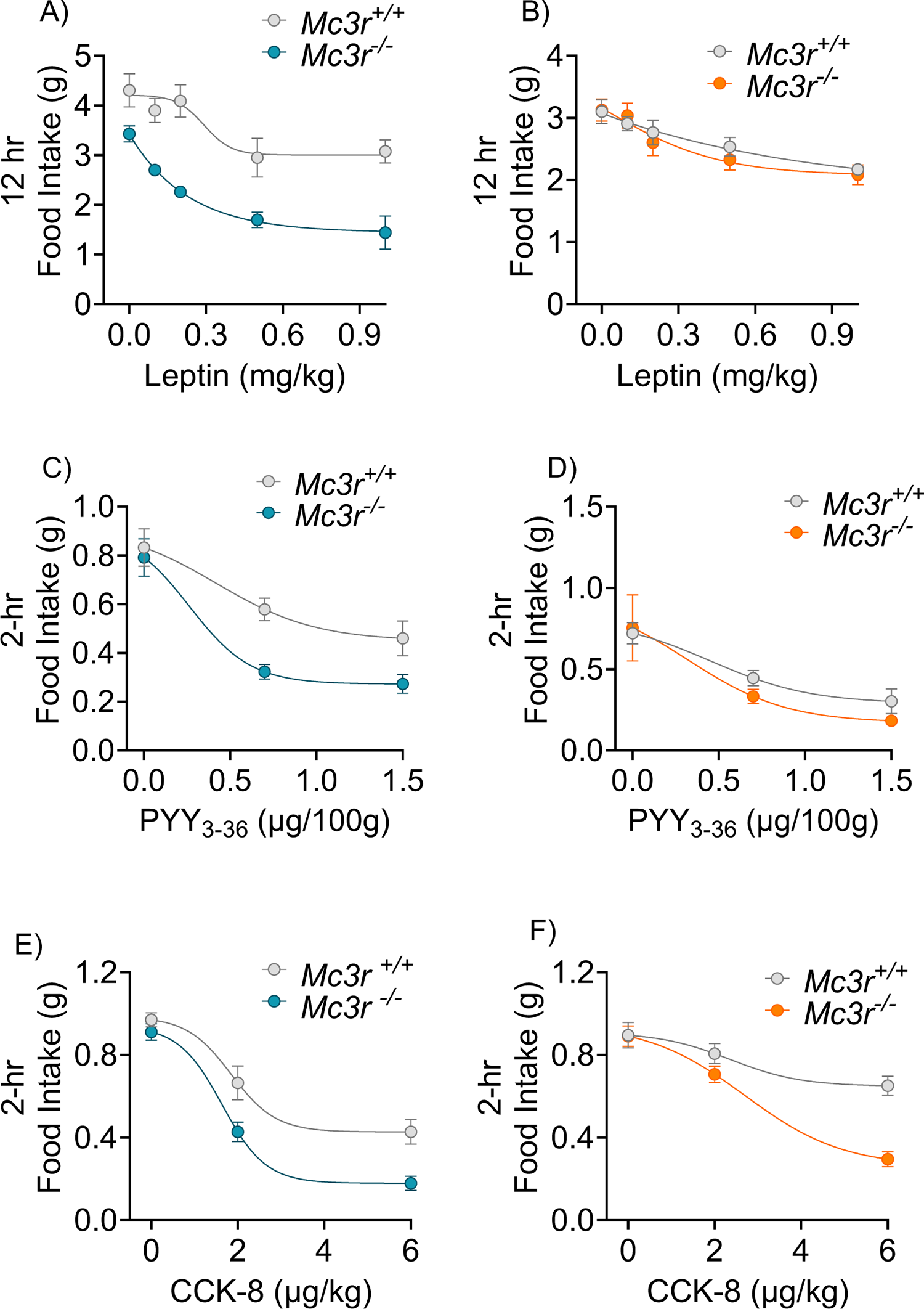
Deletion of MC3R results in generalized enhanced sensitivity to anorectic hormones. Nocturnal feeding in response to leptin responses (0.1 – 1 mg/kg) in *Mc3r ^+/+^* and *Mc3r^-/-^* at 12 hours post-injection (vehicle, n = 11, n =6/group) in *Mc3r^+/+^* and *Mc3r^-/-^* (A) male (B) female mice. Acute dark-phase feeding after administration of PYY_3-36_ (n= 7/group) and CCK ((n= 7-8/group) in *Mc3r^+/+^* and *Mc3r^-/-^* (C, E) male (F, D) female mice. Means ± SEM are shown. Statistical analysis was done with 2-way ANOVA with repeated measures (A-K) with Tukey’s post-hoc.

### Relative contributions of the MC3R and MC4R to liraglutide sensitivity

Earlier research showed that MC3R presynaptically releases GABA from AgRP neurons that synapse onto downstream MC4R neurons, highlighting that MC3R is a negative regulator of melanocortin-4 receptor-expressing (MC4R) neurons^9^. Furthermore, it has been reported that co-administration of the GLP1R-analog, liraglutide, and the MC4R agonist setmelanotide has an additive effect on weight loss^31^. Hence, we tested the hypothesis that the administration of an MC4R agonist might also increase sensitivity to liraglutide-induced inhibition of food intake and weight loss (Figure 4A-D). Peripheral administration of the MC4R peptide agonist CTX-1211 also increased the sensitivity to liraglutide-induced inhibition of food intake and weight loss in WT male mice 24 hours after treatment (Figure 4A-B). Sensitization of feeding behavior was less evident in female mice (Figure 4C), but liraglutide-induced weight loss was clearly sensitized by CTX-1211 (Figure 4D). Next, we tested if MC3R deletion would further enhance sensitization of liraglutide action by the MC4R agonist marketed to treat certain forms of genetic obesity, setmelanotide (Imcivree)^32^. Vehicle, setmelanotide (1.5mg/kg, ip.), liraglutide (0.2mg/kg, ip.), or both setmelanotide and liraglutide were administered to WT and *Mc3r ^-/-^* mice, and 24-hr food intake and changes in body weight were measured (Figure 4E-F). Both setmelanotide and liraglutide decreased food intake and promoted weight loss in WT mice, the magnitude of which was increased for both agents in *Mc3r^-/-^* mice. Notably, the anorectic and weight loss responses of the co-administration of setmelanotide and liraglutide were yet further increased in *Mc3r^-/-^* mice compared to the WT mice (Fig 4E-F).

**Figure 4:**
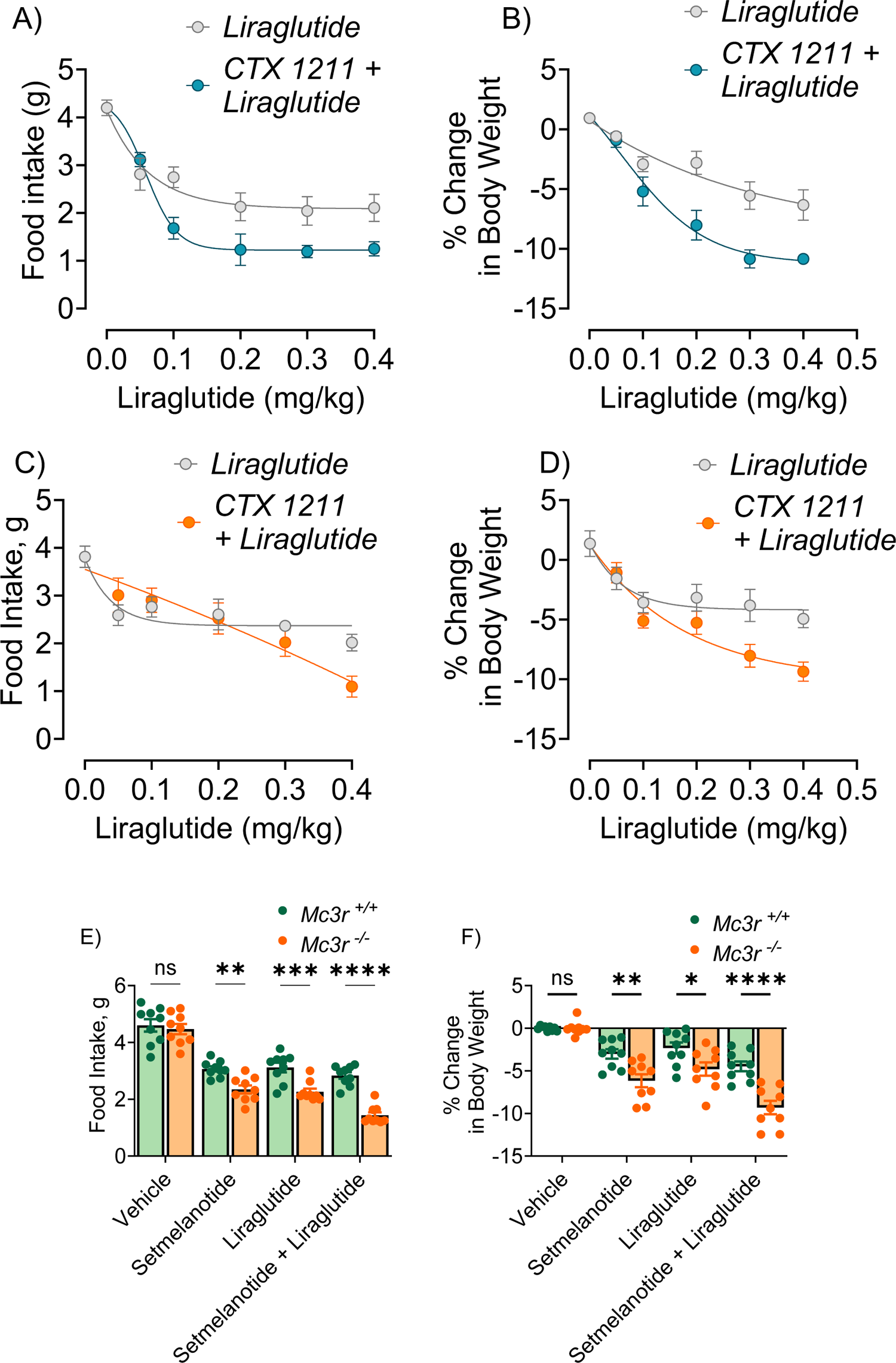
MC3R deletion enhances the ability of an MC4R agonist to increase sensitivity to liraglutide. (A, C) 24-hr food intake and (B, D) body weight changes in response to CTX-1211 (2mg/kg, i.p., n =8/group), liraglutide (0.05 mg/kg – 0.4 mg/kg, sc, n = 8) in male and female mice. (E) 24-hr food intake and (F) body weight changes in response to setmelanotide (1.5 mg/kg, i.p., n = 9), liraglutide (0.2mg/kg, sc, n= 9), liraglutide and setmelanotide (n = 9), and vehicle (n= 9) in *Mc3r^+/+^* and *Mc3r^-/-^* mice. Means ± SEM are shown. Statistical analysis was done with 2-way ANOVA with repeated measures, *p < 0.05, *p <0.01, ****p <0.001.

### Circuit mechanisms underlying anorectic hypersensitivity mediated by MC3R inhibition

Previous studies have demonstrated that much of the physiological function of the MC3R derives from the expression of the receptor in AgRP neurons. The MC3R is expressed in nearly all AgRP neurons^10^, which project widely throughout the CNS^11^. For example, the defective fasting-induced activation of AgRP neurons essential for the stimulation of food intake, observed in the global *Mc3r^-/-^* mouse, is also seen in mice with deletion of the *Mc3r* exclusively from the AgRP neurons^13^. Thus, the *Agrp-iCre;Mc3r^fl/fl^* was used to study the role of MC3R expression in the AgRP neurons in the hypersensitivity to anorectic hormones described here. Male *Agrp-iCre; Mc3r^fl/fl^* mice were more responsive to the anorectic effects of a single dose of 1mg/kg (ip) of leptin than control *Agrp-iCre* mice (Fig. 5A-B). Male and female *Agrp-iCre*; *Mc3r^fl/fl^* mice exhibited greater inhibition of feeding and greater weight loss, measured 24-hr following a 0.1mg/kg (sc) dose of liraglutide (Figure 5C-F). These data suggest that hypersensitization to both leptin and liraglutide can be recapitulated by loss of MC3R from AgRP neurons alone. Finally, we examined the role of the POMC gene in liraglutide action, encoding the MC3R and MC4R MSH agonists. Liraglutide directly activates POMC neurons and also enhances presynaptic inputs to POMC neurons^33^. Both liraglutide-mediated inhibition of food intake and weight loss were significantly blunted in male mice with a neural-specific deletion of the POMC gene; no effect was seen in female mice (Figure 6A-D). While these data do not examine the role of POMC in the hypersensitization to liraglutide by MC3R ablation/antagonism, they do implicate an important role for melanocortin signaling in liraglutide action.

**Figure 5.**
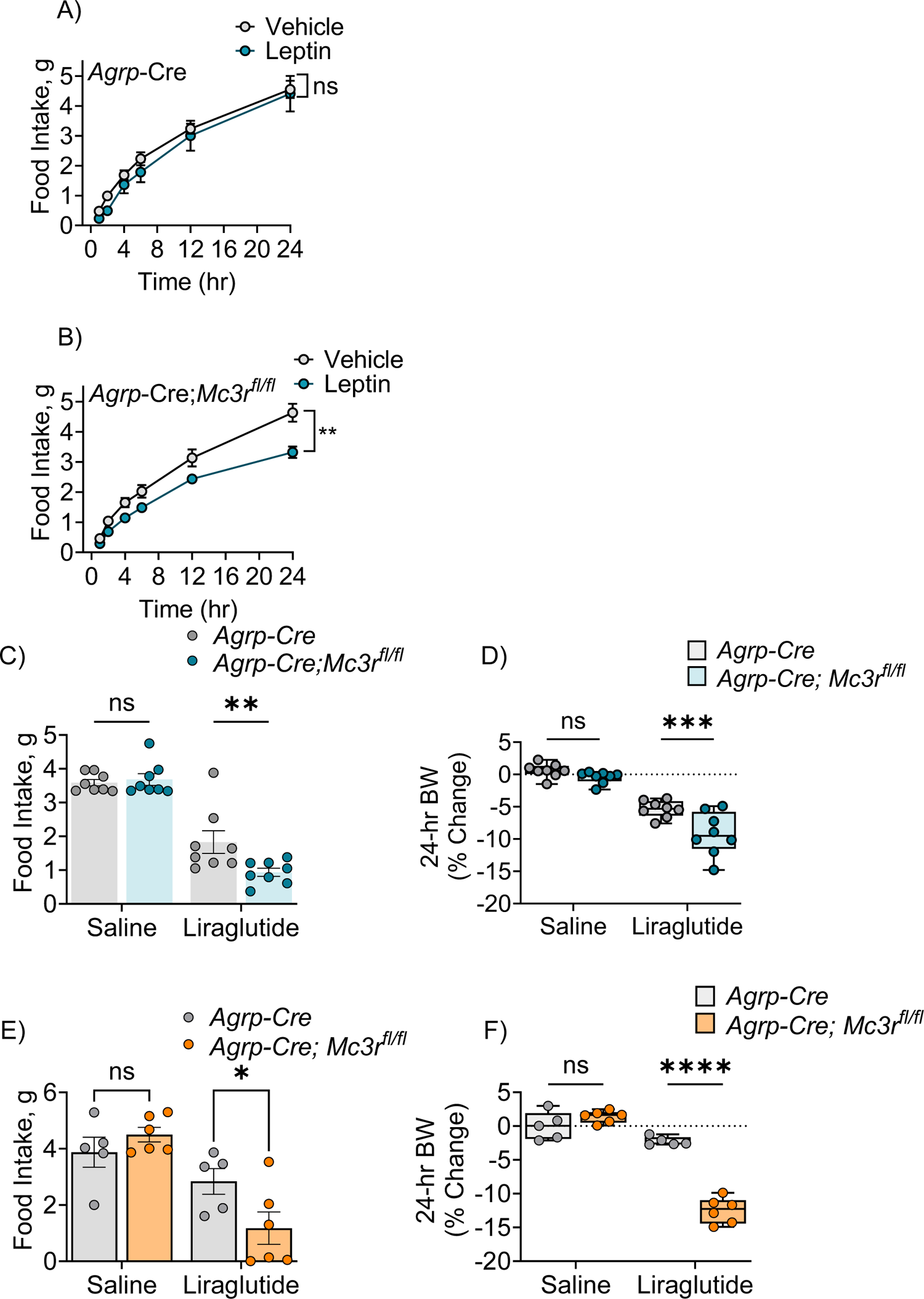
Specific deletion of MC3R in AgRP neurons results in increased responsiveness to liraglutide and leptin. (A) Time course feeding in *Agrp-iCre* and (B) *Agrp:iCre;Mc3r^fl/fl^* in response to 1mg/kg leptin (ip) (n= 8/group). 24-hr food intake and change in body weight in (C, D) males and (E, F) in females in response to liraglutide (0.1 mg/kg, sc) (n =8/group/males, n =4-6/group/females). Means ± SEM are shown. Statistical analysis was done with 2-way ANOVA with repeated measures (A, B) with Tukey’s post hoc, student t-test (C-F), *p < 0.05, **p <0.01, ****p <0.001.

**Figure 6.**
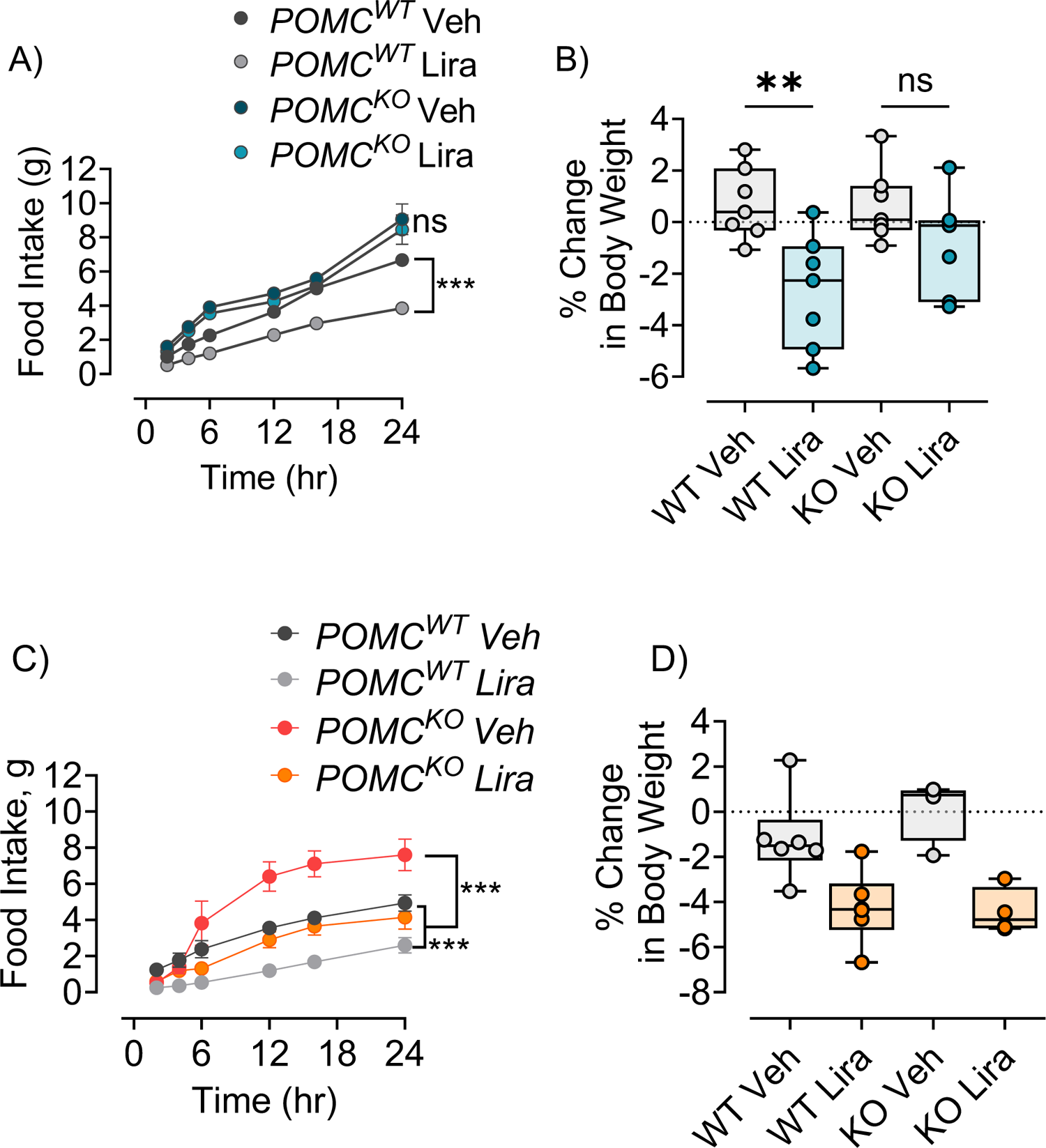
Genetic deletion of the neuronal POMC gene abolishes the response to liraglutide in males. (A, C) 24-hour food intake and percent body weight change 24-h post-injection of liraglutide in (A, B) males (0.2mg/kg, sc, n= 7/group) and (C, D) females (0.2mg/kg, sc, n=4/group). Data is expressed as mean ± SEM. Statistical analysis was done with repeated 2-way ANOVA with Tukey post hoc analysis. *p < 0.05, **p <0.01, ***p <0.001.

## Discussion

Previous studies have shown that the *Mc3r^-/-^* mouse exhibits modest obesity^14^, yet increased anorexia and weight loss in response to various challenges relative to WT littermates. For example, the *Mc3r^-/-^*was demonstrated to exhibit greater anorexia and weight loss in response to LPS treatment or IL-1β treatment and greater loss of both fat mass and lean mass in response to implantation of Lewis lung carcinoma cells^6^. More recently, we demonstrated that the *Mc3r^-/-^*mouse exhibits greater anorexia and weight loss than WT mice in response to behavioral challenges, including restraint and social isolation^10^. Further, we demonstrated that the *Mc3r^-/-^* mouse exhibits a greater extent and duration of anorexia in response to a single dose of the MC4R agonist LY2112688^9^. Here, we sought to examine this phenomenon further and determine its potential utility in the pharmacotherapy of dietary obesity. First, careful dose-response curves demonstrated that *Mc3r^-/-^* mouse exhibited increased sensitivity to the anorectic and weight loss activities of the GLP1 agonists. In striking contrast, the *Mc3r^-/-^* mouse did not show increased sensitivity to the liraglutide-induced incretin effect or malaise, as indicated by a conditioned taste aversion assay. We have demonstrated that assays of Fos immunohistochemistry, used as an indicator of neuronal activation, identified a liraglutide-induced increase in Fos signal in multiple hypothalamic feeding centers, but no increase in a region mediating emesis, the area postrema, in *Mc3r^-/-^* vs WT mice. Since mice don’t exhibit measurable emesis, it will be important to assess if MC3R antagonism can blunt the emesis induced by the GLP1 drugs in a susceptible species or if comparable weight loss can be obtained at lower GLP1 drug concentrations by MC3R antagonist treatment to avoid pharmacotherapy-induced side effects.

Importantly, the hypersensitization to GLP1 drugs could also be demonstrated using the MC3R antagonist C11 in both acute (24-hr) and chronic (3 days) treatment models. Thus, we expect that most of the anorectic hypersensitization phenomena do not result from developmental consequences of MC3R deletion and may ultimately be replicated pharmacologically. Unfortunately, the C11 molecule is difficult to synthesize, and has limited brain penetrance, thus requiring icv administration. To our knowledge, a potent, receptor-subtype specific MC3R antagonist with brain penetrance has yet to be available.

To determine if the hypersensitivity phenomenon was restricted to GLP1 analogs, we next tested the responsiveness of the *Mc3r^-/-^* mouse to both the slow-acting adipostatic hormone leptin and the rapid, acute-acting satiety factors PYY_3-36_ and CCK. The *Mc3r^-/-^* mouse is obese and hyperleptinemic and thus should exhibit leptin resistance. Remarkably, the *Mc3r^-/-^* mouse showed increased sensitivity to leptin relative to WT littermates, as assayed by 24-hour food intake and weight loss. Moreover, pharmacological inhibition of MC3R also increased leptin action by decreasing food intake and promoting weight loss in males and females (data not shown). Additional work will be needed to understand how MC3R antagonism disrupts the well-characterized phenomenon of leptin resistance. Interestingly, the *Mc3r^-/-^* mouse was also hypersensitive to the acute inhibition of feeding by the satiety factors PYY_3-36_ and CCK.

We have demonstrated that MC3R acts postsynaptically on AgRP neurons to negatively regulate MC4R neurons in the PVH^9^ and POMC neurons in the arcuate nucleus^12^. However, these same AgRP and POMC neurons project throughout the CNS to a wide variety of centers known to regulate feeding^34–39^ and to behavioral sites known to be suppressed in response to hunger and weight loss^40–42^. Thus, the data here support the hypothesis that MC3R provides negative feedback affecting circuits controlling a wide variety of hormonal and behavioral inputs to feeding and energy homeostasis. The studies shown here suggest that much of the hypersensitization phenomenon may be mediated by the disruption of MC3R signaling in AgRP neurons alone. Our studies show that MC3R antagonism may also enhance the weight loss activity of the MC4R agonist drug setmelanotide. Whether this is due to the added effects of GABA and NPY released onto MC4R neurons in addition to MC4R agonism by setmelanotide or to action on secondary sites of MC3R expression outside of AgRP neurons remains to be determined. The primary melanocortinergic AgRP, POMC neurons, and MC4R neurons are likely to play a role in the anorexigenic hypersensitivity resulting from loss of MC3R signaling. Neural deletion of POMC significantly blunts liraglutide-induced anorexia and weight loss in male mice. However, this does not directly address the hypersensitization to anorexigenic hormones seen after genetic deletion or pharmacological antagonism of the MC3R.

Rupp et al. demonstrated that neurons in the dorsomedial hypothalamus (DMH) exhibit sensitivity to both leptin and liraglutide^43^. They further found that the reactivation of *glp1r*, specifically in neurons expressing the leptin receptor, was adequate for liraglutide to effectively suppress food intake. Our observation of pronounced Fos activation in the medial basal hypothalamus (MBH) of *Mc3r^-/-^*after liraglutide treatment suggests the presence of MC3R-regulated neurons in the MBH that may be responsive to anorexigenic agents such as liraglutide and leptin. Further, significant sexual dimorphism of expression of the MC3R has been reported^8–11^, and some sexually dimorphic responses to anorexigenic agents were reported here as well. Additional work will need to be completed to determine the specific circuit mechanisms underlying the role of the MC3R in sensitization to both behavioral challenges reported previously and the wide variety of anorexigenic hormones and drugs reported here.

In conclusion, our results indicate that the loss or pharmacological inhibition of MC3R can reliably hypersensitive animals to GLP1R agonists without promoting malaise or peripheral incretin effects. Furthermore, loss of MC3R promotes generalized hypersensitivity to multiple anorexigenic agents, including CCK and leptin, short- and long-term satiety peptides, respectively. It can also, in tandem, promote the synergism of various compounds, such as in the case of MC4R and GLP1R agonists, representing the two best pharmacotherapies available for obesity and other metabolic disorders.

## Experimental Methods

### Animals models

All procedures involving animals were approved by the Institutional Animal Care and Use Committee of the University of Michigan following the American Veterinary Medical Association guidelines. Experiments were performed in C57BL/6J (wild-type, WT) purchased from The Jackson Laboratory, transgenic *Mc3r* wild-type (*Mc3r^+/+^)* and knockout (*Mc3r^-/-^)* (bred in-house), and neural-specific POMC knockout (nPOMCKO). nPOMCKO mice were generously provided by Dr. Malcom Low (University of Michigan School of Medicine). Mice were maintained on a reverse 12-h:12-h light-dark schedule and given ad libitum access to chow and water unless otherwise specified. Both sexes were used in these experiments and are shown as such in the results.

### Peptides and drugs

Liraglutide (0.05 – 0.4 mg/kg, sc, Tocris), semaglutide (1 – 100 nmol/kg, sc, Cayman Chemicals), CCK octapeptide (sulfated) (2 and 4 µg/kg, i.p., Tocris) and PYY3-36 were dissolved in a USP Grade sterile phosphate-buffered saline solution (PBS), and working concentrations were made from stock on the day of experiment. Leptin (0.1 – 1 mg/kg, *ip*., Golden West BioSolutions) was dissolved in sterile isotonic saline, and working concentrations were made on the day of the experiment. Tirzepatide hydrochloride (1-5nmol/kg, s.c., MedChem Express) was dissolved in DMSO to create stock solutions, and working concentrations were made in PBS. Setmelanotide (1.5mg/kg, i.p., Rhythm Pharmaceuticals) was dissolved in saline, and working concentrations were made on the day of the experiment. Compound 11 (Singh et al., 2013) (0.5-1nmol, *icv*.) was dissolved in DMSO. Compound CTX-1211 (2 mg/kg, *ip*) is an MC4R agonist peptide (EC_50_ = 3.8×10^-10^M) that was generously provided by Courage Therapeutics Inc. and dissolved in saline. Fresh concentrations were made on the day of the experiment.

### Acute and chronic nighttime feeding

All mice (9-14 weeks old) were single-housed in individual cages, acclimated to injection, and handled daily for 5-7 days before experimentation. Mice were randomly assigned to groups on the first day of the experiment. The number of mice used in each experiment is explained in the figure legends. The same mice were used in a crossover manner for the dose-response experiments. There were at least 4 days between each dosage. All compounds were administered 20-30 minutes before the onset of the dark, active cycle. Food was manually weighted according to experimental time points. For chronic experiments, mice received vehicle injection for at least 4 days before the start of the experiment. Compounds/peptides were injected in a volume between 150-200μl/injection. For the CCK and PYY3-336 experiments, pre-weighed chow pellets were measured before and after the acute feeding test.

### Surgery and cannula placement

C57BL/6J mice (7-8 weeks) were housed in groups of 3 with *ad libitum* access to standard chow and water before surgery. On the day of surgery, mice were anesthetized with 3-4% (v/v) isoflurane before being placed in a stereotaxic surgical frame (Kopf) and then maintained at 1.5-2% (v/v) isoflurane for the rest of the surgery. Cannulae were implanted as previously described (Sweeney et al., 2021). Guide Cannulae (Plastics One, Roanoke, VA) targeted the right lateral ventricle using the following flat skull coordinates (−0.460 mm posterior to Bregma, 1.00 lateral to the midline, and −2.20 ventral to the surface of the skull). The cannula was secured to the skull with dental cement. Following recovery, mice received an intracerebroventricular injection (*icv*) of 20 ng angiotensin II (Sigma-Aldridge) to confirm cannula placement by angiotensin-induced water intake. Cannula placement was also confirmed postmortem under a microscope.

### Saccharin-conditioned taste aversion

Age-matched WT and *Mc3r^-/-^* were handled and individually housed for 1 week with *ad libitum* access to standard chow and water. A 2-bottle test was used to administer the saccharin solution and water to mice. For the bottles, custom-made bottles (25 mL serological pipette) were fitted with rubber stoppers on one end and fabricated sipper tubes with a hole diameter of 3.175 mm (about 0.12 in) on the other end. A silicone tubing was used to seal the sipper tube and the bottle. Mice were acclimated to the 2-bottle choice and the timing of the presentation. The bottles were placed in their home cage for two days, 4-6 hours after the dark cycle, while mice received an intraperitoneal (*ip*) injection of sterile saline. Bottles were measured to ensure mice were equally drinking from the two bottles. Following acclimation, mice were conditioned for 2 days, and water was removed from the mice. On conditioning days, one group of mice received 1 mL of intraoral application of 0.1% saccharin made in water (conditioned stimulus, CS) and then immediately paired with injections (*ip*) of either 150 mM LiCl to cause gastric malaise (unconditioned stimulus, US), 0.05 mg/kg liraglutide, or 150 mM NaCl (saline) as a control solution. Mice were then given access to the saccharin solution and water in the 2-bottle tests for an additional 90 minutes. On the two testing days, mice were given access to the 2-bottle test during the typical 90-minute access; one bottle had water, and the bottle had saccharin solution. The bottles (left vs. right) were switched at the 45-minute mark to account for potential sidedness. An empty cage with a 2-bottle test was set up to account for possible spillage and was thus subtracted from each intake. Fluid intake was recorded at the end of the 90-minute test.

### Immunofluorescence staining

*Mice* were handled and injected with 150 μl sterile saline (*sc)*daily for 4 days before the experiment to minimize background neuronal activation (fos) due to stress. Immunofluorescence experiments were done to mimic the first 90 minutes of the acute experiments; thus, mice were injected with the compounds at the start of the dark to simulate the drugs’ anorectic effects. Mice received an injection of 0.1mg/kg liraglutide (*sc*) or vehicle (PBS). After 90 mins, mice were given a lethal dose of anesthesia and transcardially perfused with 4% paraformaldehyde in 0.1 M phosphate buffer (pH 7.4). Brains were removed and postfixed in the same fixative overnight. Tissues were rinsed in PBS and cryoprotected in 30% sucrose in PBS until the tissue sank. Brains were embedded in OCT (Tissue-Tek), frozen in a dry ice ethanol bath, and stored at −80 °C. Hypothalamic and brainstem sections were cut on a cryostat at 50 μm thick. The tissues were immediately washed with PBS, rocking slowly at room temperature (RT) to warm. Free-floating sections were blocked for 1 h with 0.3% triton and 5% normal serum in PBS. Primary cFos antibody (1:2000; synaptic systems 226 004) was diluted in the blocking buffer.

Sections were incubated with the primary antibody overnight, rocking at RT, rinsed 3 times in PBS, and incubated with secondary antibody in blocking buffer (Alexa conjugated antibodies, Thermo Fisher Scientific, 1:500) for 2 h at RT. Sections were rinsed 3 times with PBS, counterstained with Hoechst 33342 (Thermo Fisher Scientific, 1/2000 in PBS-0.05% Tween 20), and rinsed 2 times with PBS. Sections were moved, and slides and coverslips were mounted with Fluoromount G (Southern Biotechnology).

### Glucose or insulin tolerance tests

Mice were fasted at the onset of the light cycle for 6 hours and randomly received either treatment (0.2mg/kg liraglutide) or vehicle (PBS). Repeated OGTT occurred 2 weeks apart to ensure proper washout and recovery. Baseline blood was collected from the tail, followed by oral administration of 0.5 - 2g/kg glucose bolus in water. Blood glucose was administered over 2 h. A similar procedure was followed for an insulin tolerance test, except mice received 0.75 IU insulin/kg, and blood glucose levels were measured over 2 h. All mice received saline injections to account for fluid loss at the end of the experiment, and food was returned to the mice.

## Author contributions

NSD, PRS, and RDC designed the research studies, (NSD, YG and YW conducted the experiments and acquired the data, NSD and RDC analyzed the data, SYW, TKS, STJ, and AKM provided reagents, and NSD and RDC wrote the manuscript.

## Acknowledgments

This work was funded by NIH RO1DK126715 (RDC) and T32 DK101357 (NSD).

