## Supplemental Figures for "Inhibition of the melanocortin-3 receptor (MC3R) causes generalized sensitization to anorectic agents"

**Supplementary data**

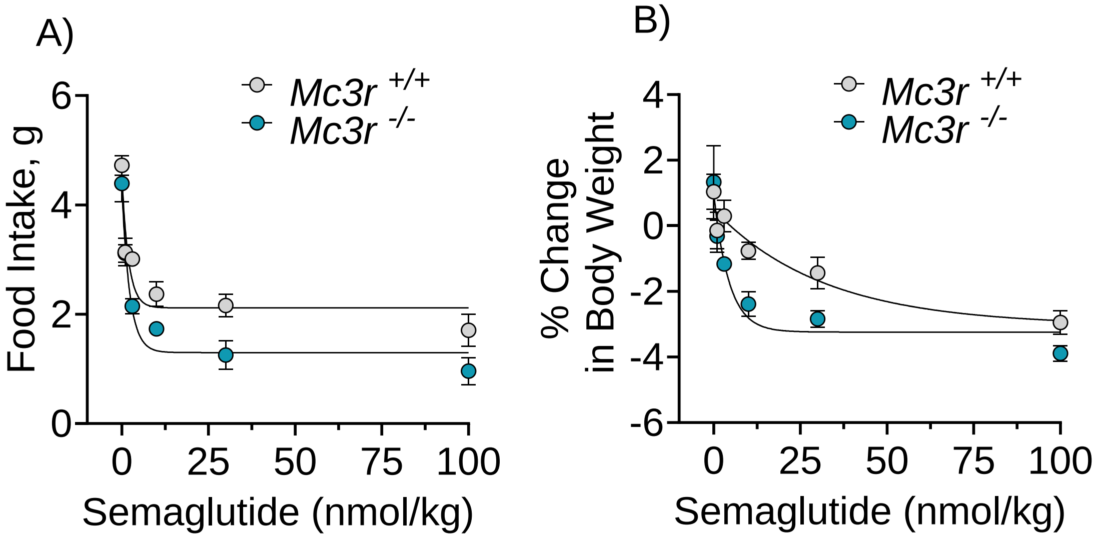


**Figure S1: Deleting MC3R increases semaglutide responsiveness.** (A-B) 24-hour feeding and percent changes in body weight in response to semaglutide (1-100nml/kg, sc, n=8/group) in *Mc3r*$^{+/+}$ and *Mc3r* $^{-/-}$ male mice. Repeated measures of two-way ANOVA were corrected for multiple comparisons using the Tukey–Kramer method for each time point, and data were fitted with four parameters: nonlinear fit.

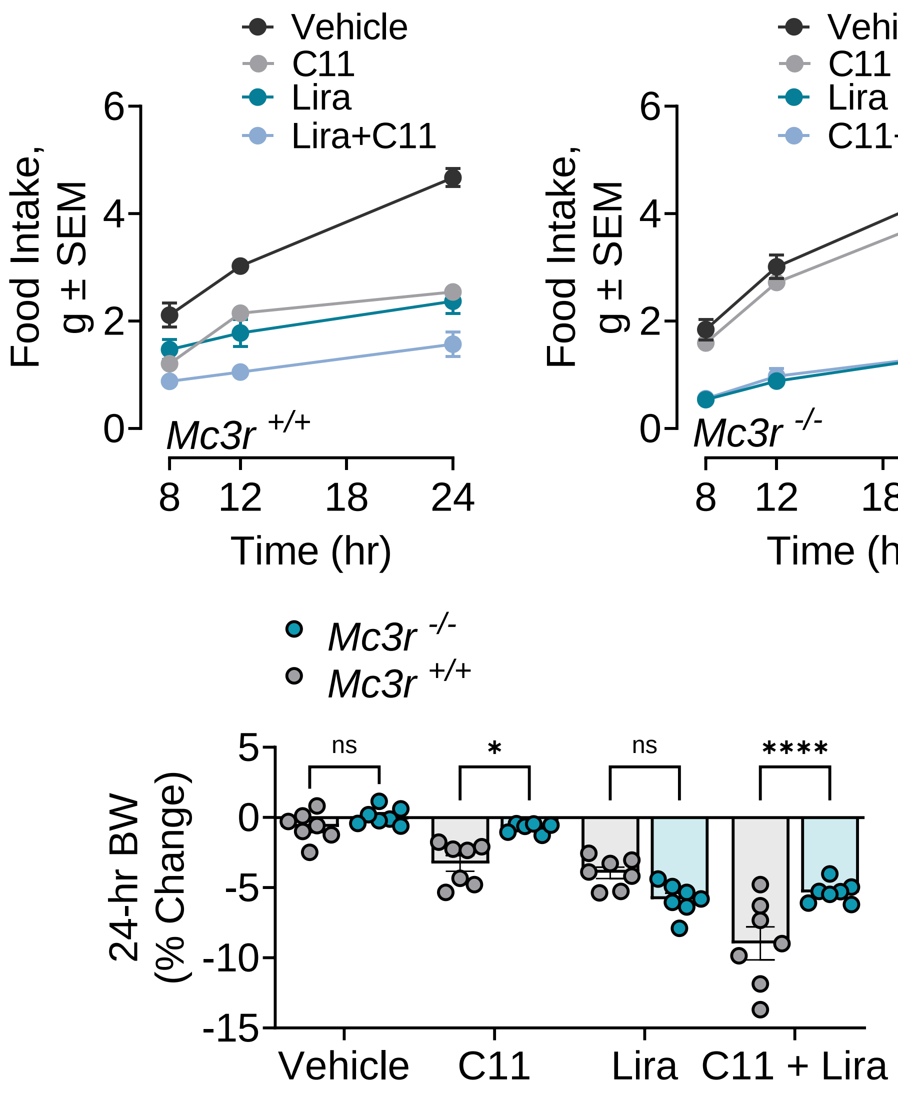


**Figure 2S: Pharmacological inhibition of MC3R increases liraglutide responsiveness.** (A-C) Time course feeding and 24-hr percent change in body weight after administration of compound 11 (1nmol/1µl, icv) or vehicle (PBS, ip, DMSO/1µl, icv). Time course (8-24 hours) feeding and 24-hour change in body weight liraglutide injection (0.2mg/kg, sc), compound 11 (C11; 1nmol/1µl, icv), a co-administration of C11 and liraglutide, and vehicle (PBS, sc; DMSO/1µl, icv of in *Mc3r* $^{+/+}$ and *Mc3r* $^{-/-}$ male mice. Statistical analysis was done with 2-way ANOVA with Tukey post hoc analysis. *$p < 0.05$, **$p < 0.01$, ****$p < 0.001$.
